# Idiosyncratic, retinotopic bias in face identification modulated by familiarity

**DOI:** 10.1101/253468

**Authors:** Matteo Visconti di Oleggio Castello, Morgan Taylor, Patrick Cavanagh, M. Ida Gobbini

## Abstract

The perception of gender and age of unfamiliar faces is reported to vary idiosyncratically across retinal locations such that, for example, the same androgynous face may appear to be male at one location but female at another. Here we test spatial heterogeneity for the recognition of the *identity* of personally familiar faces in human participants. We found idiosyncratic biases that were stable within participants and that varied more across locations for low as compared to high familiar faces. These data suggest that like face gender and age, face identity is processed, in part, by independent populations of neurons monitoring restricted spatial regions and that the recognition responses vary for the same face across these different locations. Moreover, repeated and varied social interactions appear to lead to adjustments of these independent face recognition neurons so that the same familiar face is eventually more likely to elicit the same recognition response across widely separated visual field locations. We provide a mechanistic account of this reduced retinotopic bias based on computational simulations.

**Significance statement:** In this work we tested spatial heterogeneity for the recognition of personally familiar faces. We found retinotopic biases that varied more across locations for low as compared to highly familiar faces. The retinotopic biases were idiosyncratic and stable within participants. Our data suggest that, like face gender and age, face identity is processed by independent populations of neurons monitoring restricted spatial regions and that recognition may vary for the same face at these different locations. Unlike previous findings, our data and computational simulation address the effects of learning and show how increased familiarity modifies the representation of face identity in face-responsive cortical areas. This new perspective has broader implications for understanding how learning optimizes visual processes for socially salient stimuli.

## Introduction

We spend most of our days interacting with acquaintances, family and close friends. Because of these repeated and protracted interactions, the representation of personally familiar faces is rich and complex, as reflected by stronger and more widespread neural activation in the distributed face processing network, as compared to responses to unfamiliar faces (Gobbini and Haxby, 2007; Taylor et al., 2009; Gobbini, 2010; Natu and O’Toole, 2011; Bobes et al., 2013; Sugiura, 2014; Ramon and Gobbini, 2017; Visconti di Oleggio Castello et al., 2017a). Differences in representations are also reflected in faster detection and more robust recognition of familiar faces (Burton et al., 1999; Gobbini et al., 2013; Ramon et al., 2015; Visconti di Oleggio Castello and Gobbini, 2015; Guntupalli and Gobbini, 2017; Visconti di Oleggio Castello et al., 2017b).

The advantage for familiar faces could originate at different stages of the face processing system. The classic psychological model by Bruce and Young (1986) posits that recognition of familiar faces occurs when the structural encoding of a perceived face matches stored representations (Bruce and Young, 1986). In this model the stored representations of familiar faces consist of “an interlinked set of expression-independent structural codes for distinct head angles, with some codes reflecting the global configuration at each angle and others representing particular distinctive features” (Bruce and Young, 1986, p. 309). Behavioral evidence supports the hypothesis that local features are processed differentially for personally familiar faces. For example, in a study of perception of gaze direction and head angle, changes in eye gaze were detected around 100ms faster in familiar than in unfamiliar faces (Visconti di Oleggio Castello and Gobbini, 2015). In another study, the advantage for personally familiar faces was maintained after face inversion, a manipulation that is generally thought to reduce holistic processing in favor of local processing (Visconti di Oleggio Castello et al., 2017b).

Taken together, these results suggest that optimized processing of personally familiar faces could rely on local features. This could be sufficient to initially drive a differential response to personally familiar faces. In a study measuring saccadic reaction time, correct and reliable saccades to familiar faces were recorded as fast as 180 ms when unfamiliar faces were distractors (Visconti di Oleggio Castello and Gobbini, 2015). In an EEG study using multivariate analyses, significant decoding of familiarity could be detected at around 140 ms from stimulus onset (Barragan-Jason et al., 2015). At such short latencies it is unlikely that a viewpoint-invariant representation of an individual face’s identity drives these differential responses. To account for facilitated, rapid detection of familiarity, we have previously hypothesized that personally familiar faces may be recognized quickly based on diagnostic, idiosyncratic features, which become highly learned through extensive personal interactions (Visconti di Oleggio Castello and Gobbini, 2015; Visconti di Oleggio Castello et al., 2017b). Detection of these features may occur early in the face-processing system, allowing an initial, fast differential processing for personally familiar faces.

Processes occurring at early stages of the visual system can show idiosyncratic retinotopic biases (Greenwood et al., 2017). Afraz et al. (2010) reported retinotopic biases for perceiving face gender and age that varied depending on stimulus location in the visual field and were specific to each subject. These results suggest that diagnostic facial features for gender and age are encoded in visual areas with limited position invariance. Neuroimaging studies have shown that face-processing areas such as OFA, pFus, and mFus have spatially restricted population receptive fields that could result in retinotopic differences (Kay et al., 2015; Silson et al., 2016; Grill-Spector et al., 2017b). In addition, local facial features activate the OFA (and the putative monkey homologue PL, see Issa and DiCarlo, 2012): responses to face parts are stronger when they are presented in typical locations (de Haas et al., 2016), and population activity in the OFA codes the position and relationship between face parts (Henriksson et al., 2015).

Here we hypothesized that detectors of diagnostic visual features that play a role in identification of familiar faces may also show idiosyncratic retinotopic biases and that these biases may be tuned by repeated interactions with personally familiar faces. Such biases may affect recognition of the identities presented in different parts of the visual field and may be modulated by the familiarity of those identities. We tested this hypothesis by presenting participants with morphed stimuli of personally familiar individuals that were briefly shown at different retinal locations. In two separate experiments we found that participants showed idiosyncratic biases for specific identities in different visual field locations, and these biases were stable on retesting after weeks. Importantly, the range of the retinal biases was inversely correlated with the reported familiarity of each target identity, suggesting that prolonged personal interactions with the target individuals reduced retinal biases.

We hypothesized that these biases could arise because neurons in face-processing areas have restricted receptive fields centered around the fovea (Afraz et al., 2010; Kay et al., 2015; Silson et al., 2016), resulting in an incomplete coverage of the visual field. Thus, identifying a particular face at different peripheral locations would rely on independent populations tuned to that face that cover a limited portion of the visual field biased toward the foveal region, leading to variations in identification across locations. To test this mechanism, we created a computational simulation in which increased familiarity with a specific identity resulted in changes of neural properties of the units responsive to that particular face. By either increasing the number of units responsive to a face or by increasing the receptive field size of those units, this simple learning mechanism accounted for the reduced biases reported in the two experiments, providing testable hypotheses for future work.

These findings support the hypothesis that asymmetries in the processing of personally familiar faces can arise at stages of the face-processing system where there is reduced position invariance and where local features are being processed, such as in OFA or perhaps even earlier. Our behavioral results show that prolonged, personal interactions can modify the neural representation of faces at this early level of processing, and our computational simulation provides a simple account of how this learning process can be implemented at the neural level.

## Materials and Methods

### Stimuli

Pictures of the faces of individuals who were personally familiar to the participants (graduate students in the same department) were taken in a photo studio room with the same lighting condition and the same camera. Images of two individuals were used for Experiment ī, and images of three individuals were used for Experiment 2. All individuals portrayed in the stimuli signed written informed consent for the use of their pictures for research and in publications.

The images were converted to grayscale, resized and centered so that the eyes were aligned in the same position for the three identities, and the background was manually removed. These operations were performed using ImageMagick and Adobe Photoshop CS4. The resulting images were matched in luminance (average pixel intensity) using the SHINE toolbox (function *lumMatch*) (Willenbockel et al., 2010) after applying an oval mask, so that only pixels belonging to the face were modified. The luminance-matched images were then used to create morph continua (between two identities in Experiment 1, see Figure 2; and among three identities in Experiment 1 see Figure 3) using Abrosoft Fantamorph (v. 5.4.7) with seven percentages of morphing: 0,17, 33, 50, 67, 83,100 (see Figures 2, 3).

### Experiment 1

#### Paradigm

The experimental paradigm was similar to that by Afraz et al., (2010). In every trial participants would see a briefly flashed image in one of eight locations at the periphery of their visual field (see Figure 1). Each image was shown for 50 ms at a distance of 7° of visual angle from the fixation point, and subtended approximately 4° X 4° of visual angle. The images could appear in one of eight locations evenly spaced by 45 angular degrees around fixation. For Experiment 1, only the morph *ab* was used (see Figure 1). Participants were required to maintain fixation on a central red dot subtending approximately 1° of visual angle.

**Figure 1.**
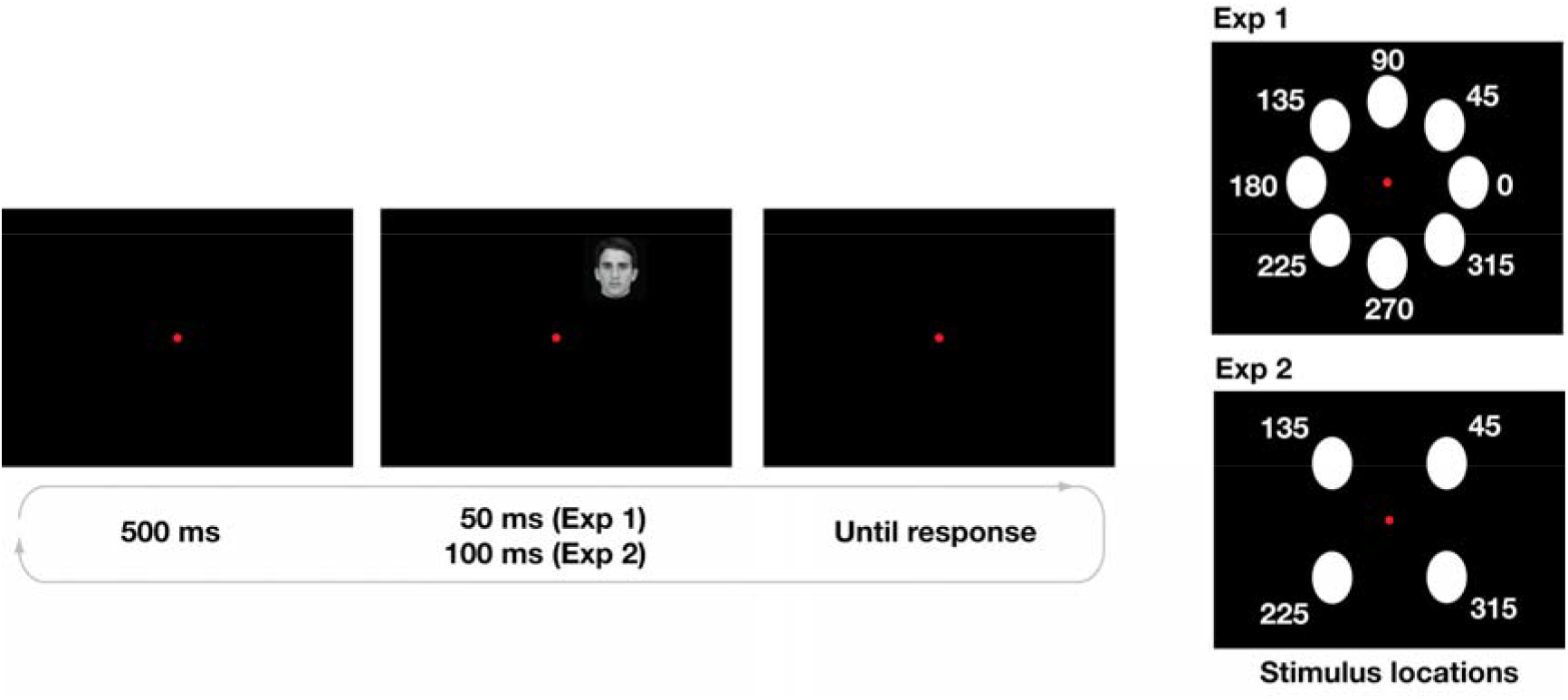
Experimental paradigm. The left panel shows an example of the experimental paradigm, while the right panel shows the locations used in Experiment ī (eight locations, top panel) and in Experiment 2 (four locations, bottom panel).

After the image disappeared, participants reported which identity they saw using the left (identity *a)* and right (identity *b)* arrow keys. There was no time limit for responding, and participants were asked to be as accurate as possible. After responding, participants had to press the spacebar key to continue to the next trial.

Participants performed five blocks containing 112 trials each, for a total of 560 trials. In each block all the images appeared twice for every angular location (8 angular locations x 7 morph percentages x 2 = 112). This provided ten data points for each percentage morphing at each location, for a total of 70 trials at each angular location.

Before the experimental session participants were shown the identities used in the experiment (corresponding to 0% and 100% morphing, see Figure 2), and practiced the task with 20 trials. These data were discarded from the analyses. Participants performed two identical experimental sessions at least four weeks apart.

**Figure 2.**
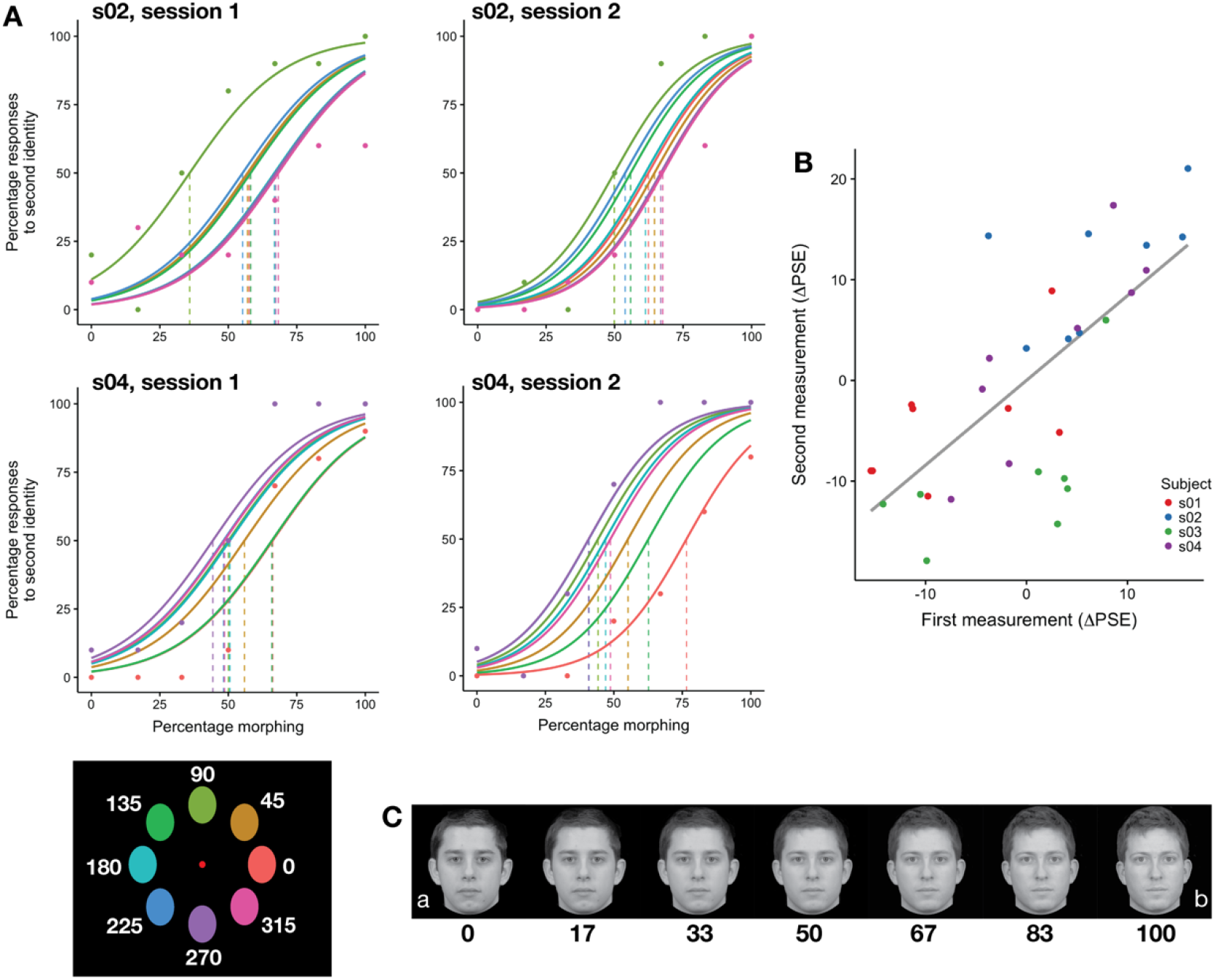
Stable and idiosyncratic biases in identification in Experiment 1. A) Psychometric fit for two subjects from both sessions. Colors indicate location (see colors in bottom left corner); actual data (points) are shown only for the extreme locations to avoid visual clutter. B) The parameter estimates across sessions (at least 33 days apart) were stable (r = 0.71 [0.47, 0.84], see Table 1). Dots represent individual parameter estimates for each location, color coded according to each subject. Correlations were performed on the data shown in this panel. C) Example morphs used in the experiment. Note that the morphs depicted here are shown for illustration only, and participants saw morphs of identities that were personally familiartothem.

Participants sat at a distance of approximately 50 cm from the screen, with their chin positioned on a chin-rest. The experiment was run using Psychtoolbox (Kleiner et al., 2007) (version 3.0.12) in MATLAB (R2014b). The screen operated at a resolution of 1920×1200 and a 60Hz refresh rate.

#### Subjects

We recruited six subjects for this experiment (three males, including one of the authors, MVdOC). The sample size for Experiment ī was not determined by formal estimates of power, and was limited by the availability of participants familiar with the stimulus identities. After the first experimental session, two participants (one male, one female) were at chance level in the task, thus only data from four subjects (two males, mean age 27.50 ± 2.08 SD) were used for the final analyses.

All subjects had normal or corrected-to-normal vision, and provided written informed consent to participate in the experiment. The study was approved by the Dartmouth College Committee for the Protection of Human Subjects.

### Experiment 2

#### Paradigm

Experiment 2 differed from Experiment 1 in the following parameters (see Figures 1, 3): 1. three morph continua (*ab, ac, bc*) instead of one; 2. images appeared in four locations (45°, 135°, 225°, 315°) instead of eight; 3. images were shown for 100 ms instead of 50 ms to make the task easier.

**Figure 3.**
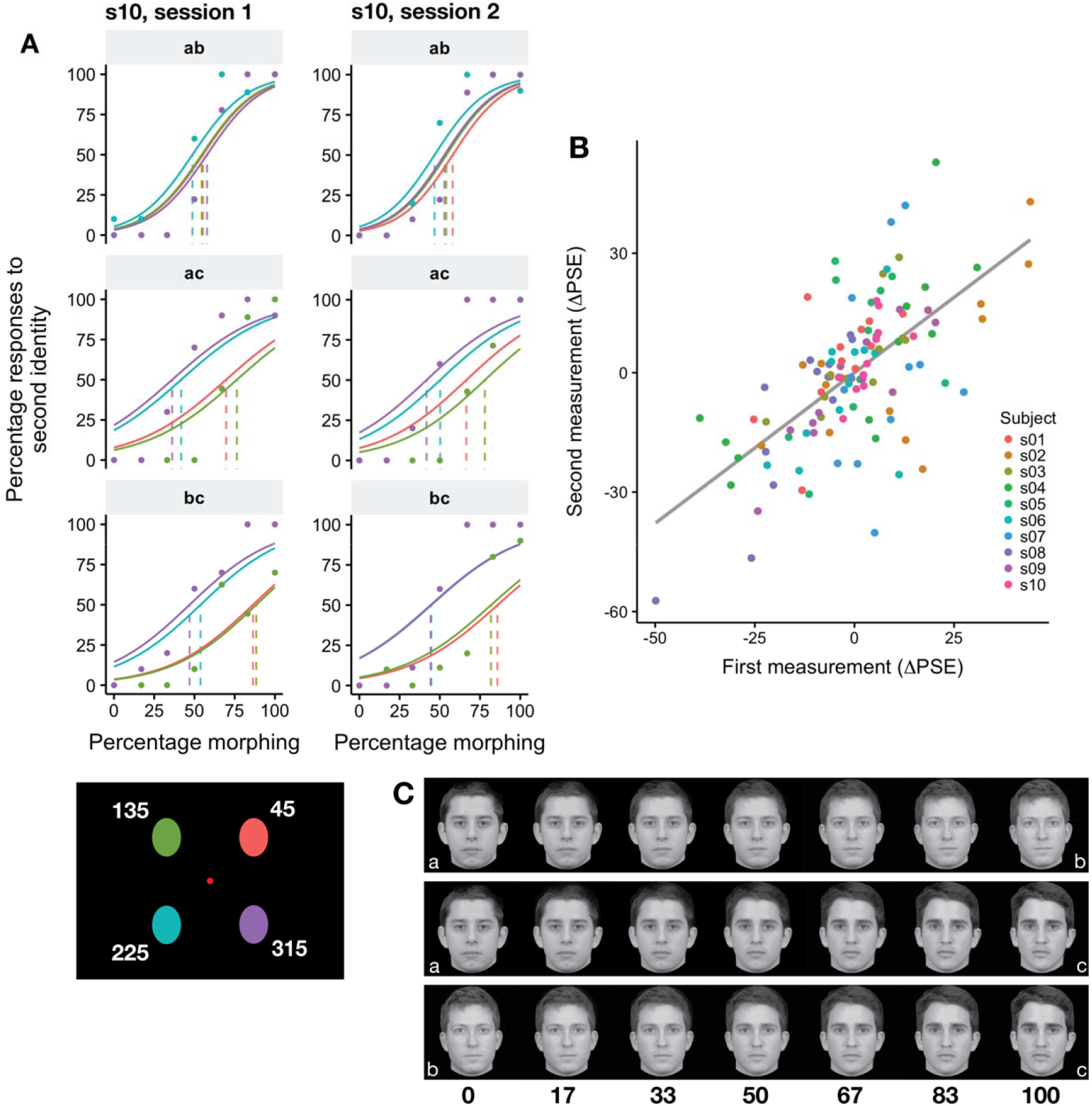
Stable and idiosyncratic biases in identification in Experiment 2. A) Psychometric fit for one subject from both sessions for each of the morphs. Colors indicate location (see colors in bottom left corner); actual data (points) are shown only for the extreme locations to avoid visual clutter. B) The parameter estimates across sessions (at least 28 days apart) were stable (r = 0.64 [0.5, 0.75], see Table 1). Dots represent individual parameter estimates for each location, color coded according to each participant. Correlations were performed on the data shown in this panel. C) Example morphs used in the experiment. Note that the morphs depicted here are shown only for illustration (participants saw morphs of identities who were personally familiar).

All other parameters were the same as in Experiment 1. Participants had to indicate which of the three identities they saw by pressing the left (identity *a*), right (identity *b*), or down (identity c) arrow keys.

Participants performed ten blocks containing 84 trials each, for a total of 840 trials. In each block all the images appeared once for every angular location (4 angular locations x 7 morph percentages x 3 morphs = 84). We used 70 data points at every angular location to fit the model for each pair of identities. Thus, we used the responses to different unmorphed images for each pair of identities, ensuring independence of the models.

Before the experimental session participants were shown the identities used in the experiment (corresponding to 0% and 100% morphing, see Figure 3), and practiced the task with 20 trials. These data were discarded from the analyses. Participants performed two experimental sessions at least four weeks apart.

#### Subjects

Ten participants (five males, mean age 27.30 ± 1.34 SD) participated in Experiment 2, five of which were recruited for Experiment 1 as well. No authors participated in Experiment 2. The sample size (*n* = 10) was determined using G*P0wer3 (Faul et al., 2007, 2009) to obtain 80% power at □ = 0.05 based on the correlation of the PSE estimates across sessions in Experiment 1, using a bivariate normal model (one-tailed).

All subjects had normal or corrected-to-normal vision, and provided written informed consent to participate in the experiment. The study was approved by the Dartmouth College Committee for the Protection of Human Subjects.

### Familiarity and contact scales

After the two experimental sessions, participants completed a questionnaire designed to assess how familiar each participant was with the identities shown in the experiment. Participants saw each target identity, and were asked to complete various scales for that identity. The questionnaire comprised the “Inclusion of the Other in the Self” scale (lOS) (Aron et al., 1992; Gächter et al., 2015), the “Subjective Closeness Inventory” (SCI) (Berscheid et al., 1989), and the “We-scale” (Cialdini et al., 1997). The lOS scale showed two circles increasingly overlapping labeled “You” and “X”, and participants were given the following instructions: *Using the figure below select which pair of circles best describes your relationship with this person. In the figure “X” serves as a placeholder for the person shown in the image at the beginning of this section, and you should think of “X” being that person. By selecting the appropriate number please indicate to what extent you and this person are connected* (Aron et al., 1992; Gächter et al., 2015). The SCI scale comprised the two following questions: *Relative to all your other relationships (both same and opposite sex) how would you characterize your relationship with the person shown at the beginning of this section?,* and *Relative to what you know about other people’s close relationships, how would you characterize your relationship with the person shown at the beginning of this section?* Participants responded with a number between one *(Not close at all)* and seven *(Very close)* (Berscheid et al., 1989). The We-scale comprised the following question: *Please select the appropriate number below to indicate to what extent you would use the term “WE” to characterize you and the person shown at the beginning of this section.*

Participants responded with a number between one *(Not at all)* and seven *(Very much so).* For each participant and each identity we created a composite “familiarity score” by averaging the scores in the three scales.

We also introduced a scale aimed at estimating the amount of interaction or contact between the participant and the target identity. The scale was based on the work by Idson and Mischel (2001), and participants were asked to respond Yes/No to the following six questions: *Have you ever seen him during a departmental event?, Have you ever seen him during a party?, Have you ever had a group lunch/dinner/drinks with him?, Have you ever had a one-on-one lunch/dinner/drinks with him?, Have you ever texted him personally (not a group message)?,* and *Have you ever emailed him personally (not a group email)?* The responses were converted to 0/1 and for each participant and for each identity we created a “contact score” by summing all the responses.

For each subject separately, to obtain a measure of familiarity and contact related to each morph, we averaged the familiarity and contact scores of each pair of identities (e.g., the familiarity score of morph *ab* was the average of the scores for identity *a* and identity *b*).

### Psychometric fit

For both experiments we fitted a group-level psychometric curve using Logit Mixed-Effect models (Moscatelli et al., 2012) as implemented in *Ime*4 (Bates et al., 2015). For each experiment and each session, we fitted a model of the form

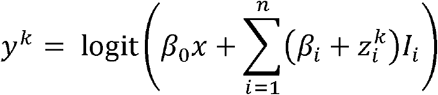

where *k* indicates the subject, *n* is the number of angular locations (*n* = *8* for the first experiment, and *n* = 4 for the second experiment), *l_i_*, is an indicator variable for the angular location, □_*i*_ are the model fixed-effects, and *z_i_* are the subject-level random-effects (random intercept). From this model, we defined for each subject the Point of Subjective Equality (PSE) as the point x such that logit(x) = 0.5, that is for each angular location

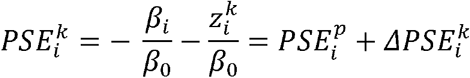

Thus, the PSE for subject *k* at angular location *i* can be decomposed in a population-level PSE and a subject-specific deviation from the population level, indicated with PSE^*p*^ and ΔPSE^*k*^ respectively.

In Experiment 2 we fitted three separate models for each of the morph continua. In addition, prior to fitting we removed all trials in which subjects mistakenly reported a third identity. For example, if an image belonging to morph *ab* was presented, and subjects responded with *c*, the trial was removed.

To quantify the bias across locations, we computed a variance score by squaring the *APSEi*, and summing them across locations, that is 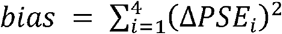. Because this quantity is proportional to the variance against 0, throughout the manuscript we refer to it as ΔPSE variance.

### Computational modeling

To account for the retinotopic biases we simulated a population of neural units activated according to the Compressive Spatial Summation model (Kay et al., 2013, 2015) and performed a model-based decoding analysis. This model was originally developed as an encoding model (Naselaris et al., 2011) to predict BOLD responses and estimate population receptive fields in visual areas and face-responsive areas such as OFA, pFus, and mFus (Kay et al., 2015). We refer to activations of neural units that can be thought as being voxels, small populations of neurons, or individual neurons.

The CSS model posits that the response of a neural unit is equal to

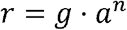

with 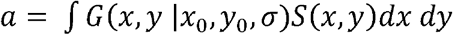, and 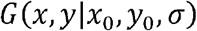 being a 2D gaussian centered at *x*_0_*y*_0_ with covariance Σ = *σI*, and *S*(*x,y*) being the stimulus converted into contrast map. The term *g* represents the gain of the response, while the power exponent *n* accounts for subadditive responses (Kay et al., 2013).

We reanalyzed the data from the fMRI experiments in Kay et al. (2015) (*pRF*-*estimation experiment* and *face-task experiment)* using the publicly available data (http://kendrickkav.net/vtcdata) and code (*http://kendrickkav.net/socmodel/*) to obtain parameter estimates for three ROIs (Inferior Occipital Gyrus, lOG—also termed OFA—mFus, and pFus). The simulation results were similar using parameter estimates from both experiments, thus we describe the procedure for the face-task experiment only because of the similarities with the behavioral experiments reported here. We refer the reader to their paper for more details on the experiments and data preprocessing. In the face-task experiment three participants saw medium-sized faces (3.2°) in 25 visual field locations (5×5 grid with 1.5° spacing), and were asked to perform a 1-back repetition detection task on face identity while fixating at the center of the screen. The resulting 25 betas were used to fit the models. As in the original paper, negative beta estimates were rectified (set to 0) and the power exponent was set to n = 0.2 and not optimized because of the reduced number of stimuli. Model fitting was performed with cross-validation. Stimuli were randomly split into ten groups, and each group was left out in turn for testing. The parameter estimates were aggregated across cross-validation runs taking the median value.

We simulated a population of *N = N_a_ + N_b_* neural units, where *N_a_* indicates the number of units selective to identity *a*, and *N_b_* indicates the number of units selective to identity *b*. For simplicity we set *N_b_* = 1 and varied *N_a_*, effectively changing the ratio of units selective to one of the two identities. We performed additional simulations increasing the total number of units and found consistent results, but here we report the simulation with *N_b_* = 1 for simplicity and consistency with the hypothesis of small neural populations responsive to specific identities. The stimuli consisted of contrast circles of diameter 4° centered at 7° from the center, and placed at an angle of 45°, 0.35°, 225°, and 315°, simulating Experiment 2. We simulated the activation of the units assuming i.i.d. random noise normally distributed with mean of 0 and standard deviation of 0.1.

Each experiment consisted of a learning phase in which we simulated the (noisy) response to the full identities *a* and *b* in each of the four locations, with 10 trials for each identity and location. We used these responses to train a Support Vector Machine (Cortes and Vapnik, 1995) with linear kernel to differentiate between the two identities based on the pattern of population responses. Then, we simulated the actual experiment by generating responses to morphed faces. For simplicity, we assumed a linear response between the amount of morphing and the population response. That is, we assumed that if a morph with *m* percentage morphing towards *b* was presented, the population response was a combination of the responses to *a* and *b*, weighted by (*1-m, m*). The amounts of morphing paralleled those used in the two experiments (0,17, 33, 50, 67, 83,100). We simulated 10 trials for each angular location and each amount of morphing, and recorded the responses of the trained decoder. These responses were used to fit a logit model similar to the model used in the main analyses (without random effects), and to estimate the Point of Subjective Equality for each angular location. The sum of these squared estimates around 50% was computed and stored.

We varied systematically the ratio *N_a_/N_b_* of units responsive to identity *a*, ranging from 1 to 9, and repeated 500 experiments for each ratio. For each experiment, parameter values (pRF location and size) were randomly sampled without replacement from the population of parameters previously estimated from the face-task experiment of Kay et al., 2015. We simulated attentional modulations by modifying the gain for the units responsive to identity *a* between 1 and 4 in 0.5 steps, and fixing the gain for identity *b* to ī. As an alternative, we simulated the effect of increases in receptive field size for the units responsive to identity *a* by increasing their receptive field size from 0% to 50% in 10% steps, while keeping the gain fixed to 1. We simulated receptive fields in this way from three face-responsive ROIs (lOG, mFus, and pFus).

### Code and data availability

Code for the analyses, raw data for both experiments, single subject results, and simulations are available at [REDACTED] as well as Extended Data.

## Results

### Experiment 1

In this experiment, participants performed a two-alternative forced-choice (AFC) task on identity discrimination. In each trial they saw a face presented for 50 ms, and were asked to indicate which of the two identities they just saw. Each face could appear in one of eight stimulus locations. Participants performed the same experiment with the same task a second time, at least 33 days after the first session (average 35 days ± 4 days standard deviation).

Participants showed stable and idiosyncratic retinal heterogeneity for identification. The PSE estimates for the two sessions were significantly correlated (see Table 1 and Figure 2B), showing stable estimates, and the within-subject correlations of ΔPSEs (see Methods) was significantly higher than the between-subject correlation (correlation difference: 0.87 [0.64, 1.10], 95% BCa confidence intervals (Efron, 1987); see Table 2), showing that the biases were idiosyncratic (see Figure 2A for example fits for two different subjects).

**Table 1.**
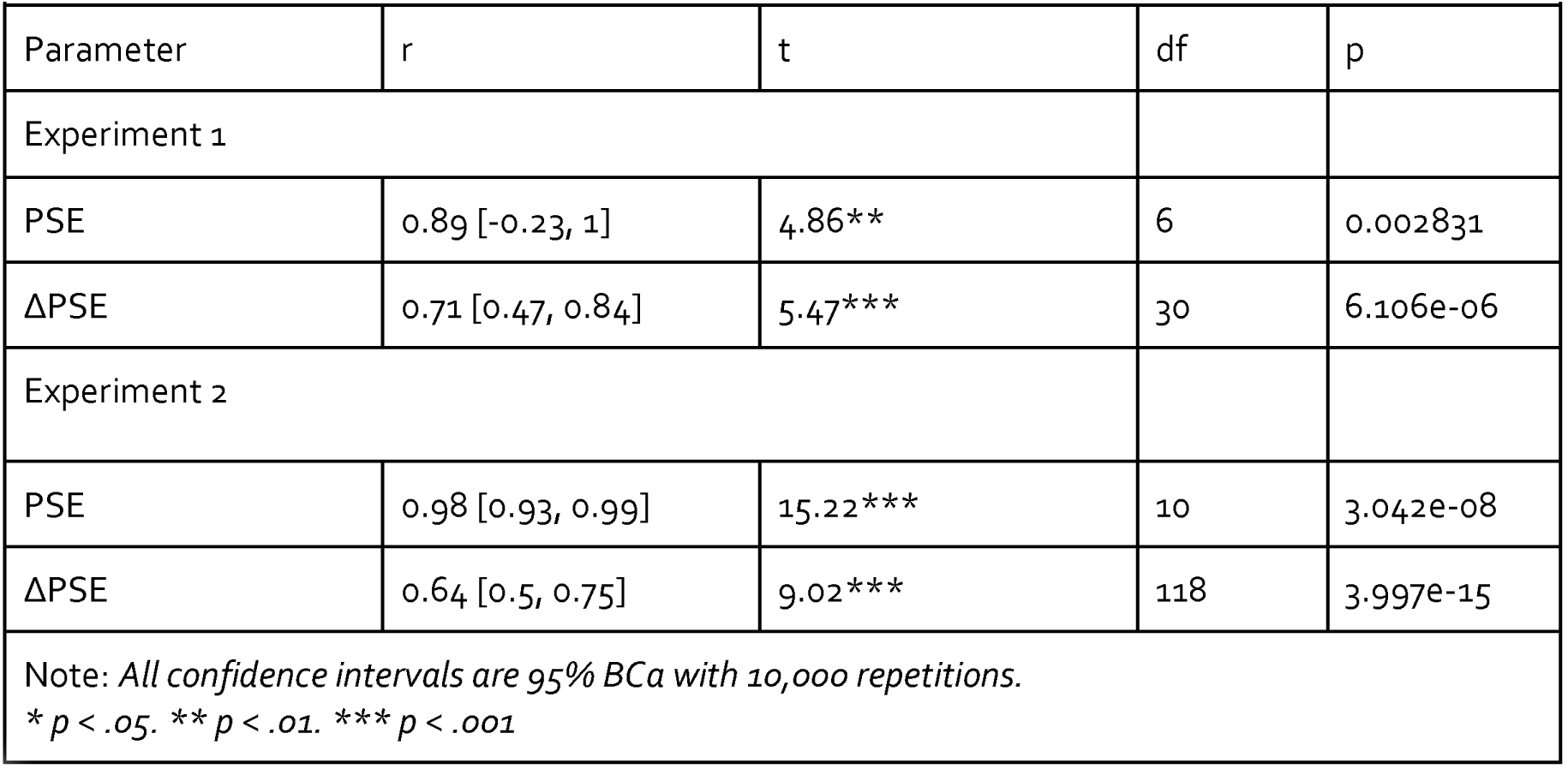
Correlation of parameter estimates across sessions for the two experiments.

**Table 2.** Comparison of within-subjects correlations of parameter estimates across sessions with between-subjects correlations.

|  |  |  |  |
| --- | --- | --- | --- |
| Morph | Within-subjects r | Between-subjects r | Difference |
| Experiment 1 |  |  |  |
| ab | 0.65<sup>†</sup> [0.57, 0.8] | -0.22 [-0.41, -0.01] | 0.87<sup>†</sup> [0.63, 1.1] |
| Experiment 2 |  |  |  |
| ab | 0.32 [-0.10, 0.62] | -0.02 [-0.15, 0.11] | 0.34 [-0.07, 0.69] |
| ac | 0.62<sup>†</sup> [0.35, 0.79] | -0.07 [-0.21, 0.08] | 0.68<sup>†</sup> [0.41, 0.92] |
| bc | 0.85<sup>†</sup> [0.61, 0.95] | -0.08 [-0.27, 0.12] | 0.92<sup>†</sup> [0.68, 1.15] |
| Note: All confidence intervals are 95% BCa with 10,000 repetitions.<br><sup>†</sup> indicates that the CIs do not contain 0. |  |  |  |

### Experiment 2

In Experiment 1 participants exhibited stable, retinotopic biases for face identification that were specific to each participant. Experiment 1, however, used only two target identities, thus it could not address the question of whether the biases were specific to target identities or to general variations in face recognition that would be the same for all target faces. For this reason we conducted a second experiment in which we increased the number of target identities. In Experiment 2, participants performed a similar task as in Experiment 1 with the following differences. First, each face was presented for 100 ms instead of 50 ms in order to make the task easier, since some participants could not perform the task in Experiment 1; second, each face could belong to one of three morphs, and participants were required to indicate which of three identities the face belonged to; third, each face could appear in four retinal locations instead of eight (see Figure 1) to maintain an appropriate duration of the experiment. Each participant performed another experimental session at least 28 days after the first session (average 33 days ± 8 days SD).

We found that participants exhibited stable biases across sessions for the three morphs (see Table 1 and Figure 3). Interestingly, within-subjects correlations were higher than between-subjects correlations for the two morphs that included the identity c (morphs *ac* and *bc*), but not for morph *ab* (see Table 2), suggesting stronger differences in spatial heterogeneity caused by identity c. To test this further, we performed a two-way ANOVA on the PSE estimates across sessions with participants and angular locations as factors. The ANOVA was run for each pair of morphs containing the same identity (e.g., for identity *a* the ANOVA was run on data from morphs *ab* and *ac*), and the PSE estimates were transformed to be with respect to the same identity (e.g., for identity *b* we considered PSE_*bc*_ and 100 – PSE_*ab*_). We found significant interactions between participants and angular locations for identity *b* (F(27, 120) = 1.77, p = 0.01947) and identity *c* (F(27, 120) = 3.34/ P = 3.229e-06), but not identity *a* (F(27,120) = 1.17, *P* = 0.2807), confirming that participants showed increased spatial heterogeneity for identities *b* and *c*. The increased spatial heterogeneity for identities *b* and *c*, but not a, can be appreciated by inspecting the ΔPSE estimates for each participant. Figure 4A shows lower bias across retinal locations for morph *ab* than the other two morphs, suggesting more similar performance across locations for morph *ab.* To investigate factors explaining the difference in performance across spatial locations between the three identities, we compared the ΔPSE estimates with the reported familiarity of the identities.

**Figure 4.**
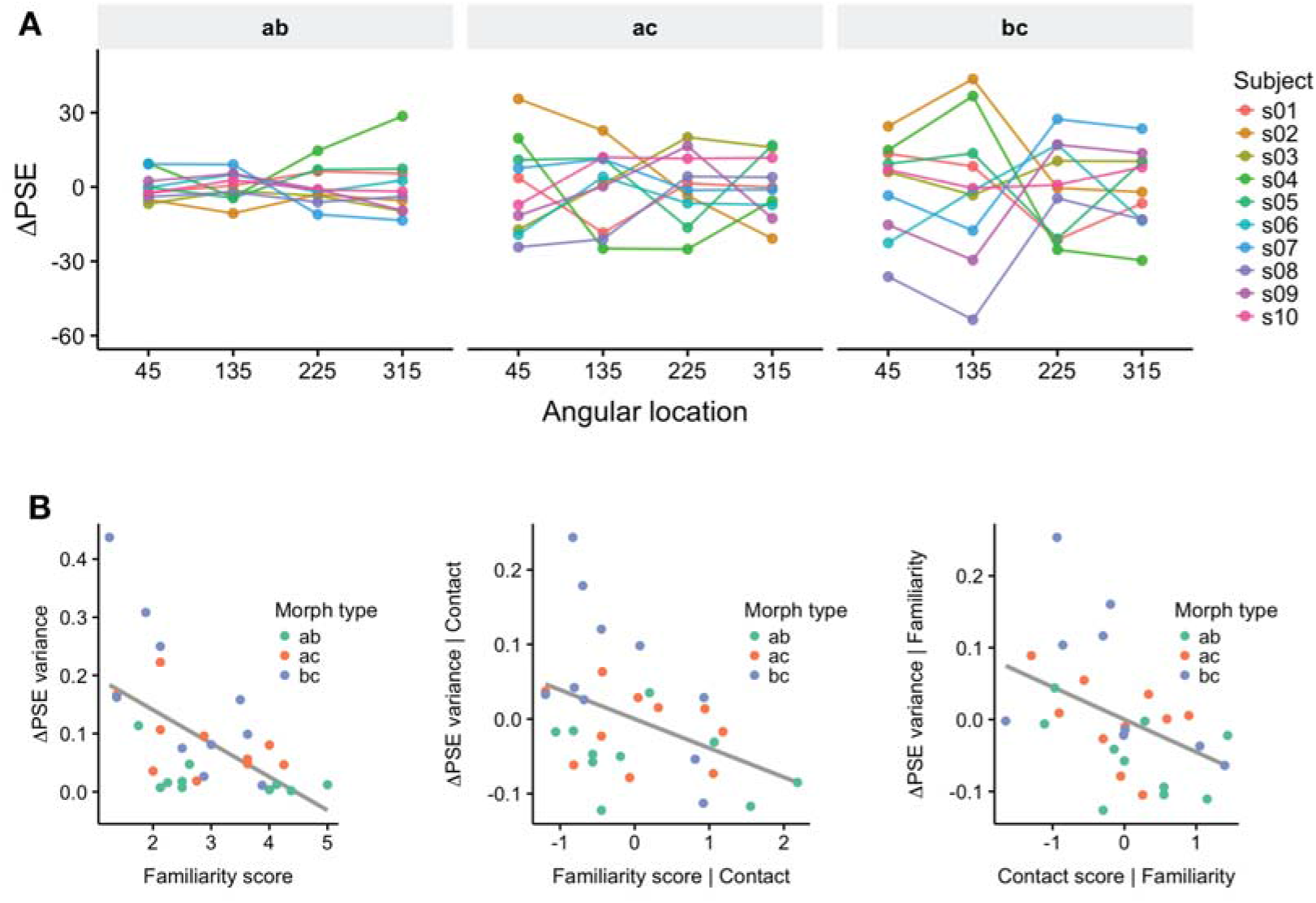
The strength of idiosyncratic biases was modulated by personal familiarity. A) Individual subjects’ ΔPSE for each morph, averaged across sessions. Note the difference in variance across locations for the three different morphs (left to right)). B) The variance across locations of ΔPSE estimates was inversely correlated with the reported familiarity of the identities (left panel; r = −0.56 [−0.71, – 0.30]), even when adjusting for the Contact score (middle panel; r_p_ = −0.42 [−0.61, – 0.16]). The right panel shows the scatterplot between the Contact score and the ΔPSE variance, adjusted for the Familiarity score, which were significantly correlated as well (r_p_ = −0.44 [−0.62, −0.17]). See Methods for definition of the Familiarity score and the Contact score. Dots represent individual participant’s data, color coded according to morph type. Correlations were performed on the data shown in these panels.

The variance of the average ΔPSE estimates across sessions for each subject was significantly correlated with the reported familiarity of the identities (r = −0.56 [−0.71, −0.30], t(28) = −3.59, p = 0.001248), showing that the strength of the retinal bias for identities was inversely modulated by personal familiarity (see Figure 4B). We estimated personal familiarity by averaging participants’ ratings of the identities on three scales (Inclusion of the Other in the Self, the We-Scale, and the Subjective Closeness Inventory, see Methods for details). The three scales were highly correlated (min correlation r = 0.89, max correlation r = 0.96).

Because the amount of personal familiarity was correlated with the amount of contact with a target identity (r = 0.45 [0.17, 0.68], t(28) = 2.65, p = 0.01304), we tested whether a linear model predicting ΔPSE with both contact and familiarity as predictors could fit the data better. Both models were significant, but the model with two predictors provided a significantly better fit (X^2^(1) = 6.30, p = 0.0121, log-likelihood ratio test), and explained more variance as indicated by higher R^2^: R^2^ = 0.45, adjusted R^2^ = 0.40 for the model with both Familiarity and Contact scores (F(2, 27) = 10.82, p = 0.0003539), 0.32, R^2^ = 0.32, adjusted R^2^ = 0.29 for the model with the Familiarity score only (F(1, 28) = 12.88, p = 0.001248). Importantly, both predictors were significant (see Table 3), indicating that familiarity modulated the variance of the ΔPSE estimates in addition to modulation based on the amount of contact with a person. After adjusting for the contact score, the variance of the ΔPSE estimates and the familiarity score were still significantly correlated (r_p_ = −0.42 [−0.61, – 0.16], t(28) = −2.42, p = 0.02235).

**Table 3.**
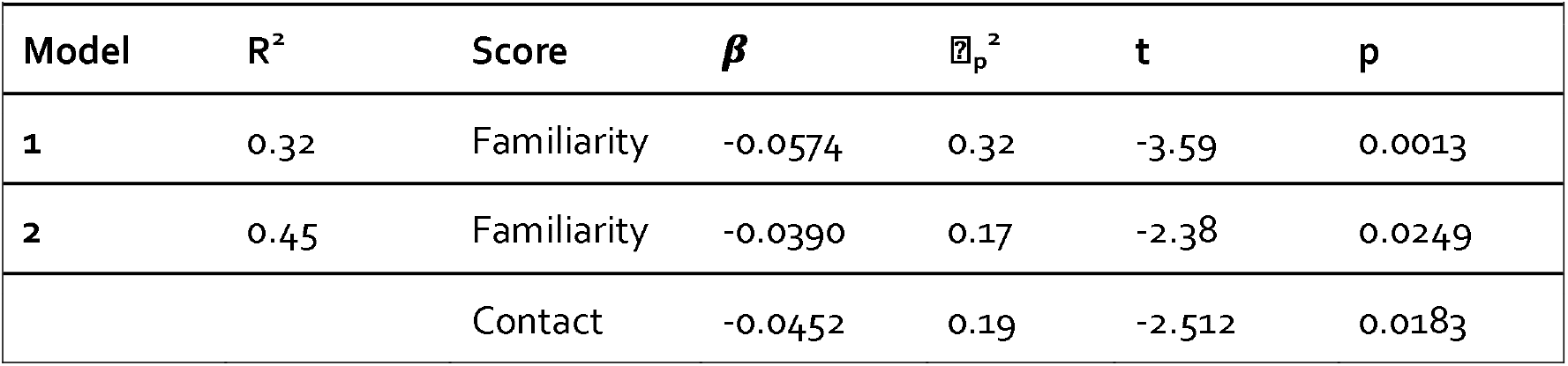
Models predicting variance of the ΔPSE estimates across locations in Experiment 2.

### Model simulation

In two behavioral experiments we found a stable, idiosyncratic bias towards specific identities that varied according to the location in which the morphed face stimuli appeared. The bias was reduced with more familiar identities, showing effects of learning. To account for this effect, we hypothesized that small populations of neurons selective to specific identities sample a limited portion of the visual field (Afraz et al., 2010). We also hypothesized that with extended interactions with a person, more neural units become selective to the facial appearance of the identity. In turn, this increases the spatial extent of the field covered by the population and thus reduces the retinotopic bias.

To quantitatively test this hypothesis, we simulated a population of neural units in lOG (OFA), pFus, and mFus activated according to the Compressive Spatial Summation model (Kay et al., 2013, 2015). The parameters of this model were estimated from the publicly available data from Kay et al. (2015). We simulated learning effects by progressively increasing the number of units selective to one of the two identities, and measuring the response of a linear decoder trained to distinguish between the two identities. As can be seen in Figure 5A, increasing the number of units reduced the overall bias (expressed as variance against 0.5 of the PSE estimates, see *Methods* for details) by increasing the spatial coverage (see Figure 5B).

**Figure 5.**
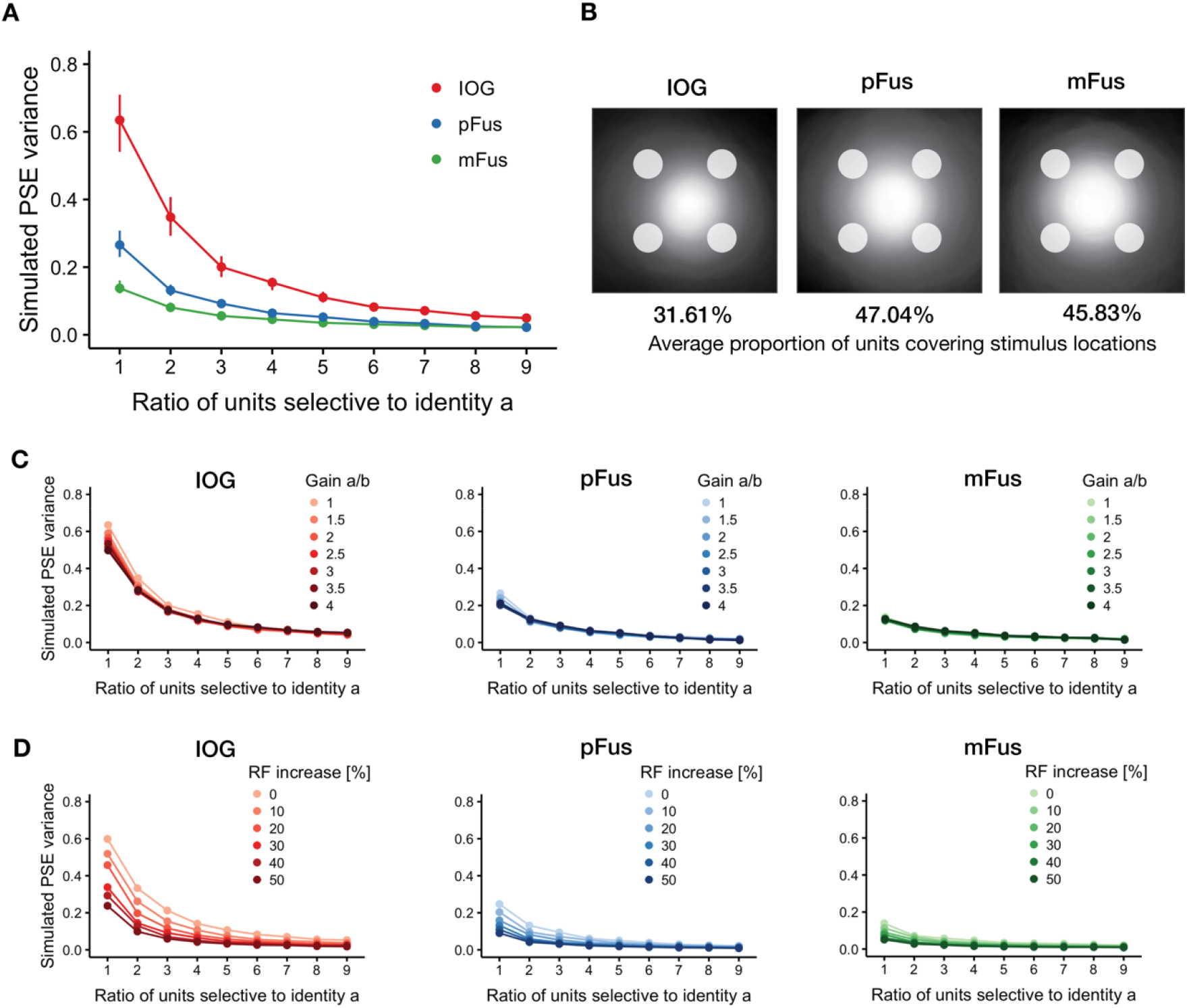
Simulating retinotopic biases and learning effects in face-responsive ROIs. We hypothesized that neural units (voxels, small populations of neurons, or individual neurons) cover a limited portion of the visual field, and that learning increases the number of neural units selective to a particular identity. A) Increasing the number of units selective to one identity reduces the retinotopic bias. Results of simulating 500 experiments by varying the ratio of neural units selective to one of two identities and fixing the gain to 1 for both identities. Dots represent median values with 95°/o bootstrapped CIs (1,000 replicates; note that for some points the CIs are too small to be seen). In all simulated ROIs the variance of the PSE around 50% decreases with increasing number of units selective to *a,* but remains larger in lOG because of its receptive field size. B) Population coverage of the units in each ROI estimated from the face-task data in Kay et al. (2015) and used in the simulations. Circles at the periphery show the simulated stimulus locations. Each image is normalized to the number of units in each ROI. Receptive fields are computed with radius 2*σ*, following the convention in Kay et al., (2015). Percentages below each image show the average proportion of units whose receptive field cover the stimulus locations. Compared to pFus and mFus, fewer units cover the stimuli in lOG resulting in a larger bias across locations. C) Increasing the gain of the response to one identity fails to reduce the retinotopic bias. D) Increasing the receptive field size of the units responsive to one identity reduces the retinotopic bias. In both C) and D) each dot represents median values of PSE variance for 500 simulated experiments. CIs are not shown to reduce visual clutter.

Interestingly, the larger bias was found within the simulated lOG. Inspecting the pRF coverage of the three ROIs revealed that the stimuli shown at 7° of eccentricity were at the border of the receptive field coverage in lOG (Figure 5B) because of the smaller RF sizes (median value across voxels of 2.98° [2.85°, 3.10°], 95% bootstrapped confidence intervals), compared to those in pFus and mFus (3.87° [3.65°, 4.05°] and 3.55° Í3.35°, 3.75°] respectively). To quantify this difference, we computed the average proportion of units covering the stimulus locations in each ROI. As predicted from the smaller RF sizes, fewer units in lOG covered the area where the stimuli were presented (31.61%) compared to pFus (47.04%) and mFus (45.83%). These results suggest that a larger retinotopic bias would be expected to originate from units in lOG.

As alternative explanations, we tested whether differences in gain or increases in RF size could reduce the bias to a similar extent as increasing the number of units. Figure 5C shows that modulating the gain failed to reduce the retinotopic bias in all simulated ROIs, while Figure 5D shows that increasing RF size of the units responsive to the more familiar identity can also reduce the retinotopic bias.

## Discussion

Afraz et al. (2010) reported spatial heterogeneity for recognition of facial attributes such as gender and age, suggesting that relatively independent neural populations tuned to facial features might sample different regions of the visual field. Prolonged social interactions with personally familiar faces lead to facilitated, prioritized processing of those faces. Here we wanted to investigate if this learning of face identity through repeated social interactions also affects these local visual processes, by measuring spatial heterogeneity of identity recognition. We measured whether face identification performance for personally familiar faces differed according to the location in the visual field where face images were presented. We found that participants exhibited idiosyncratic, retinotopic biases for different face identities that were stable across experimental sessions. Importantly, the variability of the retinotopic bias was reduced with increased familiarity with the target identities. These data support the hypothesis that familiarity modulates processes in visual areas with limited position invariance (Visconti di Oleggio Castello et al., 2017a).

These results extend the reports of spatial heterogeneity in visual processing to face identification. Similar biases exist for high-level judgments such as face gender and age (Afraz et al., 2010), as well as shape discrimination (Afraz et al., 2010), crowding, and saccadic precision (Greenwood et al., 2017). Afraz et al. (2010) suggested that neurons in IT exhibit biases that are dependent on retinal location because their receptive field sizes are not large enough to provide complete translational invariance, and stimuli in different locations will activate a limited group of neurons. In this work, we show that these perceptual biases for face processing not only exist for gender and age judgments (Afraz et al., 2010), but also for face identification and that these biases are affected by learning.

### Location-dependent coding in face-responsive areas

Neurons in temporal cortex involved in object recognition are widely thought to be invariant to object translation, that is their response to an object will not be modulated by the location of the object in the visual field (Riesenhuber and Poggio, 1999; Hung et al., 2005). However, evidence suggests that location information is preserved in activity of neurons throughout temporal cortex (Kravitz et al., 2008; Hong et al., 2016). Location information can be encoded as a retinotopic map, such as in early visual cortex, where neighboring neurons are selective to locations that are neighboring in the visual field. In the absence of a clear cortical retinotopic map, location information can still be preserved at the level of population responses (Schwarzlose et al., 2008; Rajimehr et al., 2014; Henriksson et al., 2015; Kay et al., 2015).

Areas of occipital and temporal cortices show responses to objects that are modulated by position (Kravitz et al., 2008, 2010; Sayres and Grill-Spector, 2008). In particular, also face-responsive areas of the ventral core system (Haxby et al., 2000; Visconti di Oleggio Castello et al., 2017a) such as OFA, pFus, and mFus show responses that are modulated by the position in which a face appears. Responses to a face are stronger in these areas when faces are presented foveally rather than peripherally (Levy et al., 2001; Hasson et al., 2002; Malach et al., 2002). In addition, early face processing areas such as PL in monkeys or OFA in humans code specific features of faces in typical locations. Neurons in PL are tuned to eyes in the contralateral hemifield, with receptive fields covering the typical location of the eyes at fixation (Issa and DiCarlo, 2012). Similarly, OFA responses to face parts are stronger when they are presented in typical locations (de Haas et al., 2016), and OFA activity codes the position and relationship between face parts (Henriksson et al., 2015).

The modulation of responses by object location in these areas seems to be driven by differences in receptive field sizes. In humans, population receptive fields (pRF) can be estimated with fMRI by modeling voxel-wise BOLD responses (Dumoulin and Wandell, 2008; Wandell and Winawer, 2011, 2015; Kay et al., 2013). These studies have shown that pRF centers are mostly located in the contralateral hemifield (Kay et al., 2015; Grill-Spector et al., 2017b), corresponding to the reported preference of these areas for faces presented contralaterally (Hemond et al., 2007). In addition, pRF sizes increase the higher in the face processing hierarchy, favoring perifoveal regions (Kay et al., 2015; Silson et al., 2016). The location-dependent coding of faces in these face-processing areas might be based on population activity, since these areas do not overlap with retinotopic maps in humans (for example, OFA does not seem to overlap with estimated retinotopic maps, Silson et al., 2016, but see Janssens et al., 2014; Rajimehr et al., 2014; Arcaro and Livingstone, 2017; Arcaro et al., 2017 for work in monkeys showing partial overlap between retinotopic maps and face patches).

### Cortical origin of idiosyncratic biases and effects of familiarity

Populations of neurons in visual areas and in temporal cortex cover limited portions of the visual field, with progressively larger receptive fields centered around perifoveal regions (Grill-Spector et al., 2017b). This property suggests that biases in high-level judgments of gender, age, and identity may be due to the variability of feature detectors that cover limited portions of the visual field (Afraz et al., 2010). While the results from our behavioral study cannot point to a precise location of the cortical origin of these biases, our computational simulation suggests that a larger bias could arise from responses in the OFA, given the estimates of receptive field size and eccentricity in this area (Kay et al., 2015; Grill-Spector et al., 2017b). We cannot exclude that this bias might originate in earlier areas of the visual processing stream.

In this work, we showed that the extent of variation in biases across retinal locations was inversely correlated with the reported familiarity with individuals, suggesting that a history of repeated interaction with a person may tune the responses of neurons to that individual in different retinal locations, generating more homogeneous responses. Repeated exposure to the faces of familiar individuals during real-life social interactions results in a detailed representation of the visual appearance of a personally familiar face. Our computational simulation suggests a simple process for augmenting and strengthening the representation of a face. Learning through social interactions might cause a greater number of neural units to become responsive to a specific identity, thus covering a larger area of the visual field and reducing the retinotopic biases. Our results showed that both ratings of familiarity and ratings of amount of contact were strong predictors for reduced retinotopic bias; however, familiarity still predicted the reduced bias when accounting for amount of contact. While additional experiments are needed to test whether pure perceptual learning is sufficient to reduce the retinotopic biases to the same extent as personal familiarity, these results suggest that repeated personal interactions can strengthen neural representations to a larger extent than mere increased frequency of exposure to a face. This idea is consistent with neuroimaging studies showing a stronger and more widespread activation for personally familiar faces compared to unfamiliar or experimentally learned faces (Gobbini and Haxby, 2006; Cloutier et al., 2011; Natu and O’Toole, 2011; Leibenluft et al., 2004; Gobbini and Haxby, 2007; Bobes et al., 2013; Ramon and Gobbini, 2017; Visconti di Oleggio Castello et al., 2017a).

### Effects of attention

Could differences in attention explain the modulation of retinotopic biases reported here? Faces, and personally familiar faces in particular, are important social stimuli whose correct detection and processing affects social behavior (Brothers, 2002; Gobbini and Haxby, 2007). Behavioral experiments from our lab have shown that personally familiar faces break through faster in a continuous flash suppression paradigm (Gobbini et al., 2013), and hold attention more strongly than unfamiliar faces do in a Posner cueing paradigm (Chauhan et al., 2017). These results show that familiar faces differ not only at the level of representations, but also in allocation of attention. At the neural level, changes in attention might be implemented as increased gain for salient stimuli or increased receptive field size (Kay et al., 2015). In an fMRI experiment Kay et al. (2015) reported that population receptive field (pRF) estimates were modulated by the type of task. Gain, eccentricity, and size of the pRFs increased during a 1-back repetition detection task on facial identity as compared to a 1-back task on digits presented foveally.

To address differences in gain in our computational simulation, we modified the relative gain of units responsive to one of the two identities and found that it did not influence the PSE bias across locations. This bias was more strongly modulated by the number of units responsive to one of the identities. On the other hand, simulating increases in receptive field size reduced the retinotopic bias almost as much as increasing the number of units. These simulations suggest two alternative, and possibly interacting, mechanisms that can reduce retinotopic biases in identification: recruitment of additional units selective to an identity or changes in RF properties. Additional experiments are needed to further characterize the differences in attention and representations that contribute to the facilitated processing of personally familiar faces.

### Implications for computational models of vision

Many computational models of biological vision posit translational invariance: neurons in IT are assumed to respond to the same extent, regardless of the object position (Riesenhuber and Poggio, 1999; Serre et al., 2007; Kravitz et al., 2008). Even the models that currently provide better fits to neural activity in IT such as hierarchical, convolutional neural networks (Yamins et al., 2014; Kriegeskorte, 2015; Yamins and DiCarlo, 2016) use weight sharing in convolutional layers to achieve position invariance (LeCun et al., 2015; Schmidhuber, 2015; Goodfellow et al., 2016). While this reduces complexity by limiting the number of parameters to be fitted, neuroimaging and behavioral experiments have shown that translational invariance in IT is preserved only for small displacements (DiCarlo and Maunsell, 2003; Kay et al., 2015; Silson et al., 2016; for a review see Kravitz et al., 2008), with varying receptive field sizes and eccentricities (Grill-Spector et al., 2017a). Our results highlight the limited position invariance for high-level judgments such as identity, and add to the known spatial heterogeneity for gender and age judgments (Afraz et al., 2010). Our results also show that a higher degree of invariance can be achieved through learning, as shown by the reduced bias for highly familiar faces. This finding highlights that to increase biological plausibility of models of vision, differences in eccentricity and receptive field size should be taken into account (Poggio et al., 2014), as well as more dynamic effects such as changes induced by learning and attention (Grill-Spector et al., 2017a).

### Conclusions

Taken together, the results reported here support our hypothesis that facilitated processing for personally familiar faces might be mediated by the development or tuning of detectors for personally familiar faces in the visual pathway in areas that still have localized analyses (Gobbini et al., 2013; Visconti di Oleggio Castello et al., 2014, 2017b; Visconti di Oleggio Castello and Ida Gobbini, 2015). The OFA might be a candidate for the cortical origin of these biases as well as for the development of detectors for diagnostic fragments. Patterns of responses in OFA (and neurons in the monkey putative homologue PL, Issa and DiCarlo, 2012) are tuned to typical locations of face fragments (Henriksson et al., 2015; de Haas et al., 2016). Population receptive fields of voxels in this region cover an area of the visual field that is large enough to integrate features of intermediate complexity at an average conversational distance (Kay et al., 2015; Grill-Spector et al., 2017b), such as combinations of eyes and eyebrows, which have been shown to be theoretically optimal and highly informative for object classification (Ullman et al., 2001, 2002; Ullman, 2007).

Future research is needed to further disambiguate differences in representations or attention that generate these biases and how learning reduces them. Nonetheless, our results suggest that prioritized processing for personally familiar faces may exist at relatively early stages of the face processing hierarchy, as shown by the local biases reported here. Learning associated with repeated personal interactions modifies the representation of these faces, suggesting that personal familiarity affects face-processing areas well after developmental critical periods (Arcaro et al., 2017; Livingstone et al., 2017). We hypothesize that these differences may be one of the mechanisms that underlies the known behavioral advantages for perception of personally familiar faces (Burton et al., 1999; Gobbini and Haxby, 2007; Gobbini, 2010; Gobbini et al., 2013; Visconti di Oleggio Castello et al., 2014, 2017b; Ramon et al., 2015; Visconti di Oleggio Castello and Gobbini, 2015; Chauhan et al., 2017; Ramon and Gobbini, 2017).

**Extended Data.** The archive contains data from both experiments, as well as the analysis scripts.

## Acknowledgments

We would like to thank Carlo Cipolli for helpful discussions. We would like to thank the Martens Family Fund and Dartmouth College for their support.

